# Neurotensin receptor 1 deletion suppresses methamphetamine self-administration and the associated reduction in dopamine cell firing

**DOI:** 10.1101/697656

**Authors:** Sergio Dominguez-Lopez, Ramaswamy Sharma, Michael J. Beckstead

## Abstract

We previously reported that pharmacological blockade of neurotensin receptors in the ventral tegmental area (VTA) decreases methamphetamine (METH) self-administration in mice. Here we explored the consequences of genetic deletion of neurotensin receptor 1 (NtsR1) in METH self-administration and VTA dopamine neuron firing activity. We implanted mice with an indwelling jugular catheter and trained them to nose-poke for intravenous infusions of METH. Mice with NtsR1 deletion (KO) acquired self-administration similar to wildtype (WT) and heterozygous (HET) littermates. However, in NtsR1 KO and HET mice, METH intake and motivated METH seeking decreased when the response requirement was increased to a fixed ratio 3 and when mice were tested on a progressive ratio protocol. After completion of METH self-administration, single cell in vivo extracellular recordings of dopamine firing activity in the VTA were obtained in anesthetized mice. In WT METH-experienced mice, dopamine cell firing frequency dramatically decreases compared to WT drug-naïve mice. NtsR1 KO and HET mice did not exhibit this decline of dopamine cell firing activity after prolonged METH self-administration. We also observed an increase in population activity following METH self-administration that was strongest in the WT group. Our results suggest a role for NtsR1 in METH-seeking behavior, and ablation of NtsR1 receptors prevents the detrimental effects of prolonged METH self-administration on VTA dopamine cell firing frequency.

## INTRODUCTION

Neurotensin is a 13 amino acid peptide that acts in the brain as a modulator of neurotransmission [1,2]. Most actions of neurotensin require the activation of specific G protein-coupled receptors, NtsR1 and NtsR2, which are widely distributed in the mammalian brain [3,4]. NtsR1 is observed on dopamine cell bodies and dendrites in the ventral tegmental area (VTA) and the substantia nigra (SN), as well as in dopamine axon terminals in the nucleus accumbens and the caudate/putamen [5,6]. NtsR1 is highly expressed in the dopamine neurons that are located in the rostral aspect of the VTA with well-defined mesolimbic projections [7,8]. Furthermore, the VTA receives dense neurotensinergic innervation from the lateral hypothalamus, the preoptic area, and the nucleus accumbens [9]. The overlap between the dopamine and the neurotensin systems in the mesolimbic pathway is a strong indication of a functional interaction controlling motivated behavior, including the reinforcing properties of psychostimulants such as methamphetamine (METH) [1,10,11].

Considerable evidence suggests that there may be synergism of the central effects of neurotensin and METH. METH self-administration increases neurotensin content in the striatum and midbrain [12–14]. While the actions of neurotensin in the midbrain are complex and involve multiple neurotransmitter systems [2], its overall effect is likely an increase in dopamine neuron activity [15–17]. Consistent with this, midbrain neurotensin promotes dopamine release in striatal terminals [18,19], and is itself reinforcing [20,21]. Indeed, we previously described that systemic administration of the selective NtsR1 agonist PD149163 transiently decreases motivation to self-administer METH in mice [22]. Furthermore, we also reported that a five-day treatment with microinfusions of the NtsR1/NtsR2 receptor antagonist SR142948A directly into the VTA acutely delays METH self-administration acquisition and produces a prolonged decrease in METH-seeking behavior [23]. We hypothesized that activation of VTA NtsR1 is involved in motivated METH-seeking behavior, since evoked dopamine efflux in nucleus accumbens by neurotensin application in the VTA is mostly suppressed in mice lacking NtsR1 but not in mice with NtsR2 deletion [24].

Acutely, METH has bidirectional, concentration-dependent effects on dopamine neuron firing activity [25]. We and others reported an excitatory effect of low concentrations of METH on midbrain dopamine neuron firing in brain slice and culture preparations [25,26]. Higher concentrations of METH promote a rise in extracellular dopamine through uptake inhibition and efflux, which subsequently inhibits dopamine neuron firing through activation of somatodendritic D2 type autoreceptors [27]. Consistent with this, acute systemic injections of METH dose-dependently decrease the firing frequency of dopamine neurons recorded in the VTA and the SN [28]. Our previous work suggests that D2 autoreceptor signaling is impaired after METH self-administration [29], and there is strong evidence that neurotensin decreases the functionality of D2 autoreceptors in dopamine cells [16,19,30,31]. While this might be expected to increase dopamine neuron firing in vivo, no study to date has explored how dopamine cell firing is affected by chronic METH self-administration.

Here we combined METH self-administration in mice with a genetic deletion of NtsR1 with in vivo single unit recordings of VTA dopamine neurons. We observed a decrease in METH self-administration and drug-seeking behavior in NtsR1 heterozygotes (HET) and knockouts (KO) versus wild type (WT) littermates. We also observed that METH self-administration increased the number of low firing rate dopamine neurons in vivo, an effect that was eliminated in mice lacking NtsR1.

## MATERIALS AND METHODS

### Animals

*Ntsr1^tm1Dgen^* mice (Jackson Stock #005826, Bar Harbor, ME) were generously provided by Dr. Gina M. Leinninger and were bred with C57Bl/6J mice, also from Jackson. A total of 36 adult male littermates were used, aged 6-8 months (KO, n= 10; HET, n=13; WT, n=13). Mice were group housed (3-5 per cage) in polycarbonate boxes with rodent bedding and shredding material and kept on a 12/12-hour reverse light-dark cycle (lights off at 0900 h) with *ad libitum* access to food and water. All procedures were approved by the Institutional Animal Care and Use Committee at the Oklahoma Medical Research Foundation.

### Genotyping

DNA from offspring was extracted from mouse ear punches using Extract-N-Amp^®^ kit (Sigma, St. Louis, MO) and genotyped using a multiplex PCR reaction; primers at 1 *μ*M final concentration used were: 5’ CTC TAA TGT GCC ACA GCT CAG AGA G 3’, 5’ CAG CAA CCT GGA CGT GAA CAC TGA C 3’ and 5’ CCA AGC GGC TTC GGC CAG TAA CGT T 3’. PCR conditions used were: 94°C for 30 sec, 60°C for 30 sec, and 72°C for 2 min, for 35 cycles followed by a final amplification step of 72°C for 2 min. Samples were run on 1.5% agarose gels using Tris-acetate buffer to differentiate WT, HET, and KO mice (Fig. 1A).

**Figure 1.**
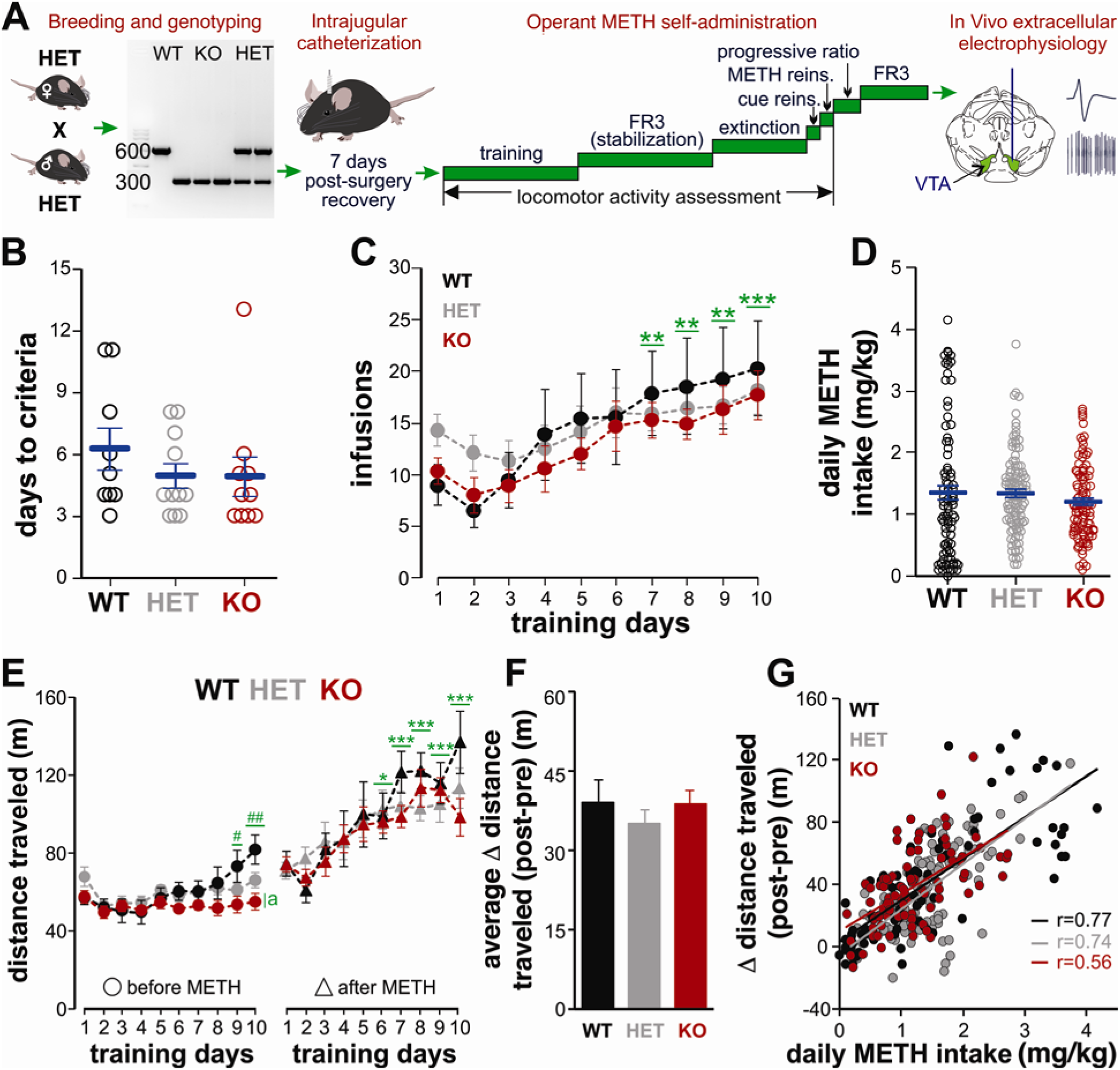
NtsR1 KO mice acquire METH self-administration normally. A) A representation of the experimental design. Neither B) The number of days required to fulfill self-administration acquisition criteria, C) the number of infusions earned on daily two-hour training sessions, or D) daily METH intake during training days was different between genotypes. E) Basal locomotor activity measured prior to METH sessions increased over time in WT, but not HET or KO mice. Locomotor activity measured after METH sessions and F) the net change in locomotion (after METH minus before METH, averaged over all 10 days) was not different between genotypes. G) The net increase in locomotion correlated positively with METH intake in all genotypes. * P<0.05, ** P<0.01, *** P<0.001 vs. day 1; # P<0.05, ## P<0.01 vs. day 2; a P<0.001 vs. WT and HET.

### Drugs

METH hydrochloride (generously provided by the NIDA drug supply program, Bethesda, MD) and chloral hydrate (Sigma-Aldrich, St Louis, MO) were dissolved in sterile saline (0.9% NaCl). Pontamine sky blue dye (Alfa Aesar, Haverhill, MA) was dissolved in sodium acetate (0.5 M, pH 7.5).

### Catheter implantation

Mice were implanted with an indwelling catheter in the right jugular vein as previously described [23]. Mice were housed individually and were allowed at least 7 days to recover from surgery. The catheter was flushed daily with 0.02 ml heparinized saline (30 IU/ml) beginning 3 to 4 days after placement.

### Operant self-administration training

Catheter patency was assessed by flushing with sterile saline before each operant session and with heparinized saline after the session finished. Two-hour operant sessions were conducted daily as previously described [23]. Two nose poke holes were located on one wall of the modular mouse operant chamber (Lafayette Instruments, Lafayette, IN), and the correct nose poke was illuminated by a dim green light located inside of the hole (see Fig. 1A). During training, responses in the correct nose poke hole were rewarded on a fixed ratio 1 schedule of reinforcement (FR1) for infusions 1-5, FR2 for infusions 6-7, and FR3 for the remainder of the session. Upon completing the response requirement, the green stimulus light was extinguished, a 15-second timeout was initiated, and METH was delivered intravenously (0.1 mg/kg/infusion, 12 μl over 2 s, accompanied by a sound stimulus of 2 kHz). METH infusion dosage was calculated based on the typical weight of an adult mouse (28 g) and corrected by actual body weight to obtain intake (mg/kg/session). Responding during the timeout was recorded but not reinforced. Self-administration was considered acquired when the number of infusions earned was ≥8 and the number of responses in the correct hole was ≥70% of the total responses for two consecutive days. 6 out of 42 total mice (4 WT and 2 HET) did not meet the acquisition criterion within 14 days and were removed from the study.

### Stabilization, extinction, and progressive ratio procedures

Following acquisition, mice were advanced to a FR3 schedule of reinforcement for 10 days (stabilization, Fig. 1A). Mice were then placed on an extinction schedule for 8 days, during which the mice were attached to the intravenous tether, but no infusions or cues (light or sound) were delivered during the 2-hour session. This was followed by one day on a FR3 cue-induced reinstatement schedule (nose pokes resulted in the sound cue, but not METH infusion). Mice were then returned to a FR3 schedule of drug self-administration. Finally, mice were placed on two days of five-hour sessions on a progressive ratio schedule. During these sessions, the number of nose pokes necessary for an infusion was gradually increased (1, 2, 4, 6, 9, …) following the procedure described by Richardson and Roberts [32].

### Locomotor activity

Before and after each 2-hour METH session, mice were placed in activity chambers with infrared photosensors (Columbus Instruments, Columbus, OH). Spontaneous locomotor activity was quantified for 15 min (Fig. 1A).

### Single cell-extracellular recordings

Animals were anesthetized with chloral hydrate (400 mg/kg, i.p.) and placed in a stereotaxic apparatus (Kopf Instruments, Tujunga, CA). A hole was drill in the skull to give access to the brain. Body temperature was maintained at 35-36.5°C using an infrared heating lamp. Single-barrel glass micropipettes with an internal diameter of 1.5 mm (Sutter Instruments, Novato, CA) were pulled to a shank length of approximately 0.6 cm in a Narishige PC-10 puller (Tokyo, Japan) and filled with 2% pontamine sky blue dye. Electrodes (5-8 MΩ) were lowered into the VTA using a hydraulic microdrive (Stoelting, Wood Dale, IL) within the stereotaxic coordinates described in the mouse brain atlas of Franklin and Paxinos [33](A-P: −2.9 to −3.6 mm from Bregma, lateral: 0.2 to 0.7 mm from the midline and ventral: 3.5 to 5.5 mm from the brain surface). One to five electrode descents were performed per mouse. The spontaneous electrical activity of single cells was recorded using an MDA-4 amplifier system (Bak Electronics, Umatilla, FL). Spike frequencies and waveforms were collected and stored on a computer using Spike 2 and a Micro1401-3 acquisition unit (CED, Cambridge, UK). Dopamine neurons were identified based on well-established electrophysiological properties: a broad action potential (> 2.5 ms), biphasic or triphasic waveform, and slow firing (0.5–10 Hz) [34,35]. Neuronal activity was measured by calculating the mean firing rate frequency, expressed as the number of spikes per second, and the number of spontaneously active neurons per electrode descent or track (neurons/track).

Additionally, the burst activity of dopamine neurons was analyzed using a Spike 2 script, based on criteria previously described [36]. A burst was defined as a train of at least two spikes with an initial interspike interval (ISI) ≤ 80 ms and a maximum ISI of 160 ms within a regular low-frequency firing pattern and decreased amplitude from the first to the last spike within the burst [34,35]. The parameters analyzed included the percentage of spikes fired in bursts, the mean burst ISI, and the inter-burst interval. On the final track for each mouse, pontamine sky blue dye was injected iontophoretically by passing a constant positive current of 20 μA for 5-10 min through the recording pipette to mark the recording site. Mice were then decapitated, and their brains extracted and placed in paraformaldehyde solution (4%) for at least two days. Subsequent localization of the labeled site was made by visual inspection of 40 μm-thick brain coronal sections.

### Statistical Analysis

Data were analyzed using SigmaStat 4.0 statistical suite (Systat Software Inc., San Jose, CA). A one-way ANOVA was used to compare days to acquisition between groups. Two-way ANOVA (repeated measures when allowed) was used to determine interactions between genotype and either daily session or schedule. For *post-hoc* comparisons after ANOVAs, Bonferroni-corrected t-tests were used. Correlation of METH intake with distance traveled was done using Pearson product moment correlation, and the correlation coefficients (r) were then compared using Fisher’s Z-transformation. All data are reported as the mean ± standard error of the mean. Statistical values of p ≤ 0.05 were considered significant.

## RESULTS

### NtsR1 deletion does not affect acquisition of METH self-administration

We first aimed to determine how deletion of NtsR1 affects the ability of mice to learn to intravenously self-administer METH. Overall no differences between genotypes were observed in the average days required to meet acquisition criteria (F_2,27_=0.74, P=0.48, Fig. 1B). No effect of genotype was observed either in the average number of correct responses (F_2,243_=0.43, P=0.65) or the number of infusions delivered (F_2,243_=0.23, P=0.8) during the first ten days of training. Independently of genotype, the number of responses (F_9,243_=9.94, P<0.001) and infusions (F_9,243_=13.08, P<0.001) increased by training day 7 (Fig. 1C), consistent with our previous observations in other strains [23]. The average intake of METH across training days was not different between genotypes (P=0.84, Fig. 1D). By the final day of training, mice in all groups exhibited high accuracy for the active nose poke hole (WT: 88.94 ± 2.5%; HET: 86.88 ± 3.9%; KO: 87.93 ± 4.0%). Our results indicate that deletion of NtsR1 does not interfere with learning of intravenous METH self-administration.

As METH is a powerful psychomotor stimulant, we also monitored spontaneous locomotion in all mice before and after METH self-administration sessions. On the first day of training, no significant differences between genotypes were observed on basal locomotor activity (F_2,27_=2.39, P=0.11). In the subsequent training days, METH-induced locomotion steadily increased (F_9,247_=10.94, P<0.001, Fig 1E) but was independent of genotype (F_2,247_=1.98, P=0.14). By training day 10, basal (pre-session) locomotion had also increased, but not in the KO mice (F_2,247_=9.4, P<0.001). Analysis of net change in locomotion (post-pre) also indicated a steady increase during training (F_9,247_=7.29, P<0.001), with no differences between genotypes (F_9,247_=0.757, P=0.47; Fig. 1F). In all groups, a positive correlation between the daily METH intake and the net increase in locomotion was detected in training sessions (WT: r=0.77, P<0.001; HET: r=0.74, P<0.001; KO: 0.56, P<0.001, Fig. 1G).

### Deletion of NtsR1 decreases METH self-administration

To evaluate the influence of NtsR1 deletion on METH reinforcement, we first increased the response requirement to FR3. On this schedule, the number of responses (F_2,264_=28.06, P<0.001), infusions (F_2,264_=27.93, P<0.001, Fig. 2A) and METH intake (F_2,264_=30.91, P<0.001, Fig. 2B, 2C) was higher in WT compared with both HET and KO mice. Mice in all groups continued to exhibit a high level of accuracy for the active nose poke hole (WT: 90.08 ± 1.2%; HET: 85.45 ± 1.0%; KO: 84.77 ± 1.2%). Similarly, WT mice displayed higher basal (pre-session) locomotion (F_2,227_=41.39; Fig. 2D), higher METH-induced (post-session) locomotion (F_2,227_=30.03, P<0.001), and a higher change (post-pre) in locomotion (F_2,227_=10.44, P<0.001, Fig. 2E). Positive correlations between METH intake and the increase in locomotion were again observed in all genotypes (WT: r=0.72, P<0.001; HET: 0.85, P<0.001; KO: r=0.637, P<0.001, Fig. 2F). This suggests that, while acquisition of METH self-administration is unaffected by deletion of NtsR1 receptors, METH intake on a FR3 schedule is reduced.

**Figure 2.**
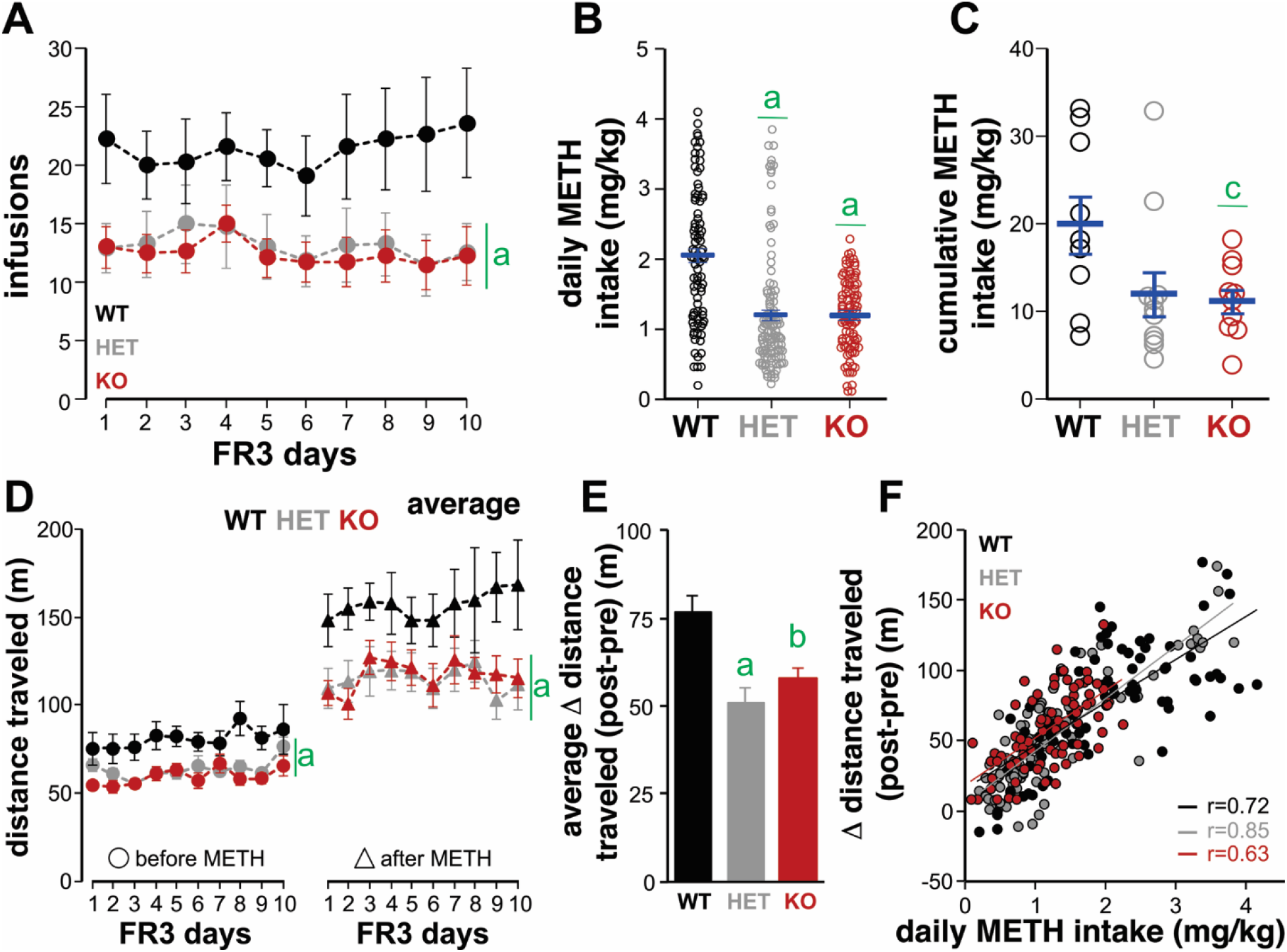
NtsR1 HET and KO mice self-administer less METH than WT mice on a FR3 schedule. Compared to WT mice, NtsR1 HET and KO mice exhibit A) a decreased number of METH infusions and B) decreased daily METH intake. C) Cumulative METH intake per mouse over the ten FR3 sessions was lower in KO mice. D) Locomotor activity measured before and after METH sessions was lower in HET and KO mice. E) The net change in locomotion after METH sessions (averaged over all 10 days) was higher in WT mice. F) In all groups, the net increase in locomotion correlated positively with daily METH intake. a P<0.001, b P<0.01 and c P<0.05 vs. WT.

### Deletion of NtsR1 decreases METH-seeking behavior

Following 10 days at FR3, we used an extinction protocol followed by cue and METH-induced reinstatement to evaluate drug-seeking behavior [23]. In the absence of both cues and METH infusions (i.e., FR ∞), the number of correct responses on the first day of extinction was lower in HET and KO mice compared with WT mice (F_16,196_=2.3, P=0.004, Fig. 3A). No differences were observed between groups in the number of extinction sessions necessary to decrease the number of responses to less than 50% (WT: 4.8 ± 0.5 days; HET: 5.2 ± 0.5 days; KO: 5.6 ± 0.7 days; F_2,27_=0.42, P=0.65). One WT and one KO mouse did not meet criteria for extinction within eight sessions and were subsequently excluded from further analysis.

**Figure 3.**
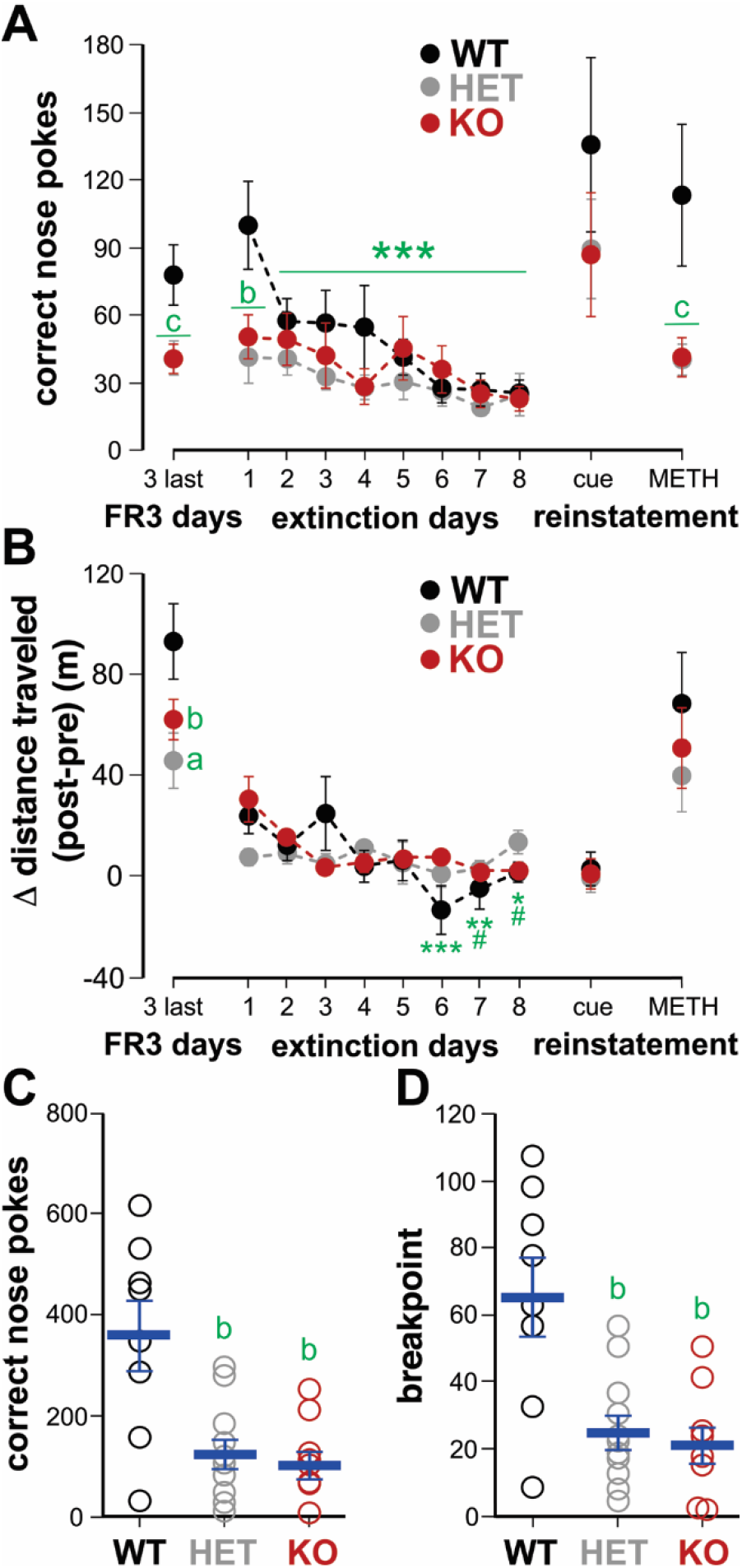
Deletion of NtsR1 decreases METH seeking behavior. A) Compared to HET and KO mice, WT mice showed greater responding on extinction day 1 and METH reinstatement, but not cue reinstatement. B) Net change in locomotion (post-pre) decreased in all groups in the absence of METH on extinction and cue reinstatement days, but was not changed on METH reinstatement day from the average of the last 3 FR3 days. In the progressive ratio protocol, C) HET and KO mice exhibited a lower number of responses and D) the breakpoint was also lower compared with WT mice. * P<0.05, ** P<0.01, *** P<0.001 vs. extinction day 1 (WT only); # P<0.05 vs. extinction day 1 (KO only); a P<0.001, b P <0.01, c P<0.05 vs. WT.

Following extinction, we reinstated the cues associated with METH infusion on a FR3 schedule, and the number of correct side nose pokes increased in all groups (F_1,25_=28.11, P<0.001). Although WT mice performed a higher number of responses (increase from last extinction day 8; WT: 499%, HET: 361%, KO: 352%), this increase between genotypes was not statistically significant (F_2,25_=0.713, P=0.5). The following day, METH was reinstated along with the cues on a FR3 schedule. In this case, WT mice exhibited a higher number of responses (F_2,25_=5.6, P=0.01, see Fig 3A), infusions (F_2,25_=6.6, P=0.005) and METH intake (F_2,25_=7.07, P=0.004) versus HET and KO groups. This indicates that all genotypes strongly associate the cues with METH infusions and that drug-seeking differences due to NtsR1 deletion are most robustly observed in the presence of METH.

Locomotor analysis showed that on extinction days, a general decrease in net locomotor activity was present (F_14,149_=1.98, P=0.023, Fig. 3B). No differences between genotypes were detected in locomotion for cue and METH reinstatement sessions. This indicates that differences between genotypes in METH-induced locomotion may be sensitive to recent METH intake history, with differences becoming evident when self-administration remains constant for several days.

Finally, to test if NtsR1 deletion affects motivation to obtain METH, we tested the mice with a protocol that systematically increases the response requirement for METH infusion [23,32]. In this progressive ratio test, KO and HET mice performed fewer nose pokes than WT mice (F_2,25_=10.3, P≤0.001, Fig. 3C), resulting in a lower number of infusions (F_2,25_=5.68, P=0.009) and lower METH intake (F_2,25_=7.16, P=0.003). Consequently, the maximum ratio completed by HET and KO mice (breakpoint) was lower than that observed in WT mice (F_2,25_=10.17, P<0.001; Fig. 3D).

### NtsR1 deletion protects against METH-induced decline in dopamine neuron activity

We next investigated the effects of NtsR1 deletion and METH self-administration on VTA dopamine neuron firing. To control for total lifetime METH intake across groups, following the progressive ratio experiment we returned mice to additional FR3 sessions. At the time of recording, the average cumulative METH intake was not significantly different between groups (WT=52.09±4.7 mg/kg; HET=60.35±6.4 mg/kg; KO=52.85±4.5 mg/kg; F_2,9_=0.675, P=0.53). We recorded 58 neurons from mice that had performed METH self-administration (WT: n=21, HET: n=8; KO: n=29) and 34 neurons from drug naïve mice (WT: n=12, HET: n=10; KO: n=12). Due to the low number of dopamine neurons recorded in METH-experienced HET mice, some comparisons were made only between KO and WT mice. Surprisingly, dopamine neuron firing rate was considerably reduced in WT mice after METH self-administration exposure (F_1,70_= 5.76, P=0.019, Fig. 4A), but this decrease was not observed in HET or KO mice.

**Figure 4.**
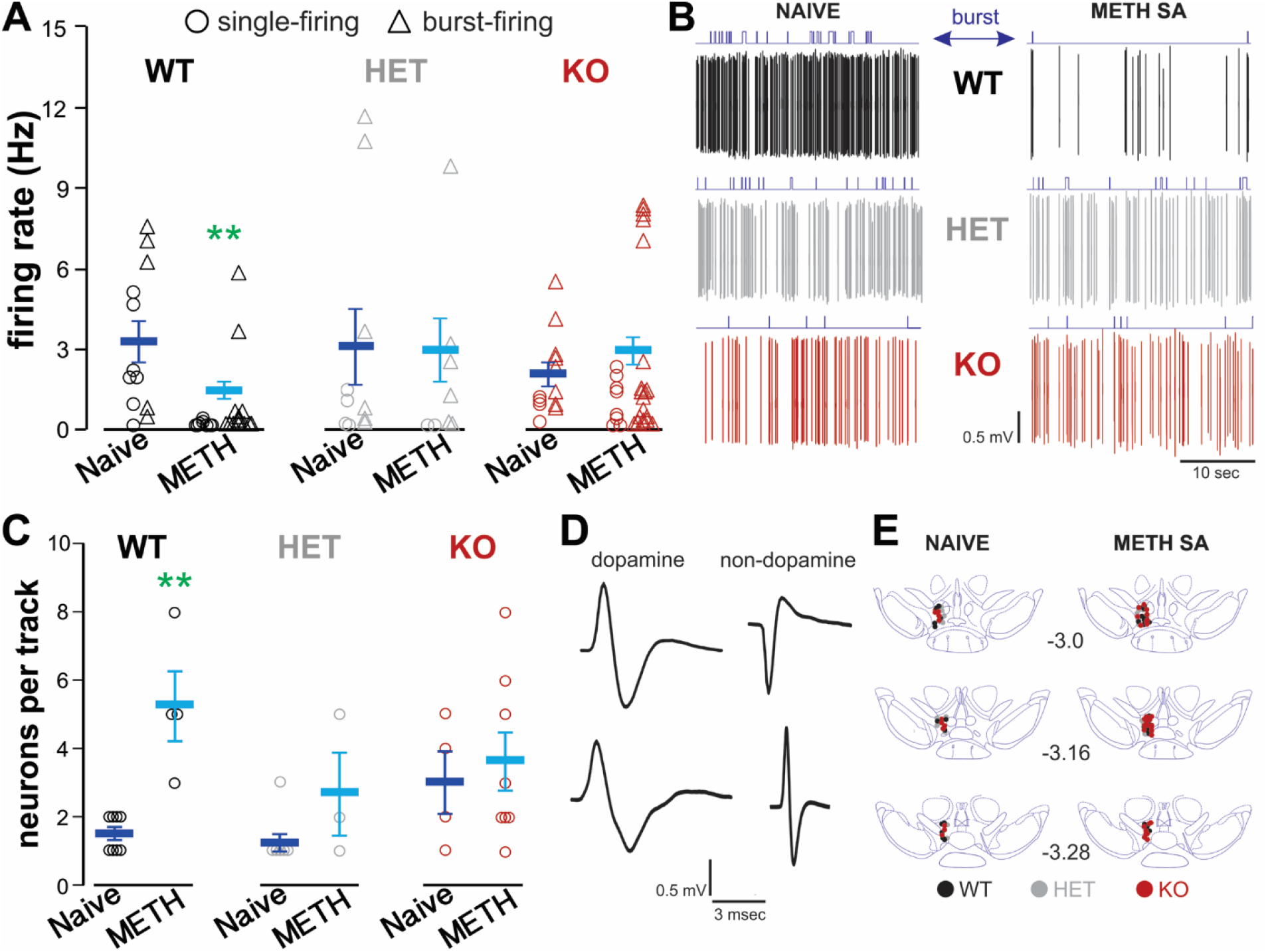
METH self-administration decreases VTA dopamine firing activity, but not in NtsR1 KO mice. A) Single-cell extracellular recordings in anesthetized mice showed a decrease in the firing rate of dopamine neurons recorded in WT mice that had performed METH self-administration (METH SA), but not in NtsR1 HET and KO mice. B) Sample recordings of dopamine neuron firing activity in all groups; burst activity is indicated in blue on top. C) The number of spontaneously active neurons recorded per electrode track increased in WT animals after METH self-administration. D) Representative traces of dopamine and non-dopamine neurons waveforms. E) Recording sites in the VTA for all groups. ** P<0.01 vs. WT naive.

Using established burst parameters [36], we did not observe significant differences in the proportion of cells firing bursts between genotypes or METH exposure conditions (Naïve: WT=41.6%, HET=60%, KO=66.6%; METH exposed: WT=61.9%, HET=75%, KO=72.4%). Both neuronal populations (single spike and burst firing) exhibited decreased firing rate after METH self-administration exposure in WT mice, but the decrease fell short of statistical significance in burst firing cells (single firing: WT: F_1,23_=8.3, P=0.008; burst firing: F_1,43_ 3.7, P=0.06, Fig. 4A). Independent of genotype, we also detected a decrease in intra-burst ISI (naïve: 41.6 ± 5.1 ms; METH exposed: 27.2±3.1 ms; F_1,43_=5.6, P=0.02) and an increase in time between bursts (naïve: 5.14±3.73 sec; METH exposed: 16.25±2.31 sec; F_1,43_=6.42, P<0.015) in neurons recorded from METH-experienced mice. To our knowledge, these data are the first evidence that prolonged METH self-administration leads to decreased firing and more sporadic burst activity in VTA dopamine neurons (Fig. 4B).

Finally, we analyzed the population activity or the number of recorded neurons that fired spontaneously per electrode descent [35]. Overall, we observed an increase in population activity in mice after METH self-administration (F_1,20_=7.66, P<0.012), an effect that was largely driven by a significant increase in the WT group (Fig. 4C). Together, our results indicate that METH self-administration produces an increase of low firing rate dopamine neurons with decreased burst capability; this effect is reduced in mice that lack of NtsR1 receptors.

## DISCUSSION

The present study tested the hypothesis that NtsR1 is involved in the development of METH-seeking behavior and determined the effects of prolonged METH self-administration on the firing activity of VTA dopamine neurons in the presence and absence of NtsR1. The results show that mice with a genetic deletion of NtsR1 exhibit normal acquisition of METH self-administration behavior, but reduced METH intake and seeking behavior when the operant requirement is increased. These observations are not related to changes in basal locomotor activity, however reduced locomotor behavior following METH sessions in NtsR1 HET and KO mice correlate well with their reduced intake of the psychomotor stimulant. We also report for the first time that prolonged METH self-administration in WT mice increases the prevalence of slow firing VTA dopamine neurons with erratic bursting. This change in firing activity was not observed in mice lacking NtsR1.

### Role of NtsR1 in METH self-administration and seeking behavior

In the VTA, neurotensin acts as a behavioral reinforcer and has been reported to both increases mesolimbic dopamine neurotransmission and enhance behavioral sensitization to psychostimulants [37–40]. These effects are thought to be substantially mediated by NtsR1, as pharmacological blockade prevents the expression of psychostimulant behavioral sensitization, but not the acute increase in behavior [37,41–44]. VTA dopamine neurons express NtsR1 [8], and psychostimulants are known to increase neurotensin input to the VTA. Even a single exposure to METH can produce an increase of neurotensin mRNA expression in nucleus accumbens neurons that project to the VTA [45,46]. This finding may account for an initial neurotensin-mediated priming to the salience of METH exposure and, as METH self-administration progresses, other brain areas are recruited. Indeed, increased neurotensin levels in the nucleus accumbens shell, dorsal striatum, SN and VTA are observed after continuous METH self-administration [12,13]. Further, on the first day of extinction following METH self-administration the midbrain exhibits a transient increase of neurotensin levels while the dorsal striatum and globus pallidus show a decrease [14]. NT has complex effects on synaptic transmission throughout the brain [2], and although the VTA is a major node for its actions on METH reinforcement other regions likely also play a role in the present observations.

The observed effects on METH reinforcement are apparently not due to changes in locomotor behavior. In agreement with others, we did not observe altered spontaneous locomotor activity in NtsR1 KO mice [47–49], although other groups have reported increased basal locomotion in more extended observation periods [50,51]. Here we observed that, even after prolonged exposure to METH self-administration, the level of basal activity before the self-administration session in NtsR1 HET and KO mice remained unchanged. We did observe an increase in WT mice that was consistent with conditioning to the operant chamber combined with the higher levels of intake for these mice. We also observed an increase in net locomotor increase following self-administration session. Interestingly, which this was repeatedly observed during stabilization at FR3 it was not seen on the METH reinstatement day following extinction, arguing against a role for a decrease in persistent behavioral sensitization in the NtsR1 HET and KO mice. Given the strong relationship in all groups between open field locomotion and METH intake it is likely that the observed effects on locomotion are simply due to differences in pharmacological exposure.

Although NtsR1 is the most prominent central neurotensin receptor [2], it is also likely that others also play a role is METH reinforcement. We previously reported that blockade of neurotensin receptors in the VTA with the NtsR1/NtsR2 antagonist SR142948A microinjected prior to the sessions on the first five days of training delays the acquisition of METH self-administration [23]. After cessation of the drug these mice acquire self-administration normally, although their METH intake and METH-seeking behavior is decreased weeks later [23]. Our current data show that deletion of NtsR1 reduces METH intake and seeking but does not affect acquisition, suggesting that VTA NtsR2 may play a specific role in the initial effects of METH. It is also interesting that the NtsR1 HET mice behaved in all experiments more similar to KO than WT mice, suggesting that even a partial reduction of NtsR1 can blunt METH reinforcement. It is also important to note that developmental effects may also play a role, given the constitutive nature of the NtsR1 KO. Experiments with more elegant approaches (such as cell type-specific NtsR1 and NtsR2 knockouts) will be required to definitively determine the role of specific receptors in specific brain regions in the effects of METH.

### Effects of METH self-administration on dopamine neuron impulse activity

In the VTA, NtsR1 is known to affect dopamine neuron activity through multiple mechanisms that are almost exclusively excitatory [2]. In dopamine neurons themselves, NtsR1 produces a depolarization through the opening of intrinsic nonselective cation channels [15,17,52]. Incredibly, at least three neurotransmitter systems that provide input to VTA dopamine neurons are also affected by neurotensin. NtsR1 receptors are responsible for an increase in long-term potentiation of NMDA receptor-mediated EPSCs in VTA dopamine neurons [11]. Neurotensin has been repeatedly linked to decreased D2 autoreceptor signaling in dopamine neurons, an effect that has been attributed to both NtsR1 [15,30,31,53] and by our lab to NtsR2 [16]. GABA input to dopamine neurons is also affected in a complex manner by both NtsR1 and NtsR2 [2,31]. Here, we report no effect of genotype on basal dopamine neuron firing in naïve mice. However, following METH self-administration we observed a dramatic decrease of dopamine cell firing and possibly burst activity. This effect was dependent on NtsR1 as it was only observed in WT mice, and was surprising given previous reports of increased dopamine neuron firing in rats that had self-administered cocaine [54,55]. Further, although data describing cells firing observed per track can be difficult to interpret, we also observed an increase in population activity following METH self-administration that was only significant in the WT group. We limited our recordings to the rostral VTA where a higher number of NtsR1 expressing dopamine neurons are observed [7,8]. However, a percentage of the recorded cells may not express NtsR1 and may have only been affected by the presence or absence of the receptor on synaptic inputs. Again, cell-type specific approaches will be necessary to shed light on which VTA neurotensin mechanisms contribute to the effects we observe following repeated METH self-administration.

## FUNDING AND DISCLOSURE

This work was supported by a National Institute on Drug Abuse grant to MJB (R01 DA032701) and funds obtained through the Presbyterian Health Foundation. The authors declare no conflict of interest.

## ACKNOWLEDGEMENTS

We want to thank Dr. Gina Leinninger for generously providing the NtsR1 KO line, Melinda West for assistance with genotyping, and Dr. Amanda Sharpe for a critical read of the manuscript.

